# DIURETIC AND ANTI-HYPERTENSIVE ACTIVITY OF *CLERODENDRUM CHINENSE* (OSBECK) MABB. AQUEOUS EXTRACT IN 8% SALT DIET INDUCED HYPERTENSIVE RATS

**DOI:** 10.1101/2023.05.09.539974

**Authors:** Joy I. Odimegwu, Tolulope F. Okanlawon, Oluwakayode Ezekiel Olatunji, Ismail Ishola

## Abstract

Diuresis refers to increase in the rate of urine flow and sodium excretion from the system via the urine. It is a necessary excretory process that may prove difficult for some disease systems e.g. enlarged prostates. Hypertension occurs as a result of systolic blood pressure higher than 140 mmHg or a diastolic blood pressure higher than 90 mmHg. It is one of the most common chronic diseases affecting more a billion people worldwide. A high dietary sodium intake is one of the factors associated with the development of hypertension. *Clerodendrum chinensis* is used by local communities in West Africa for its diuretic and anti-hypertensive properties. We analyse the phytochemical properties of the mixed leaf, root and stem aqueous extract of the plant and investigate its anti-hypertensive and diuretic activity in Sodium chloride diet induced hypertensive rats. The anti-hypertensive effect of extract at different concentrations (100, 200 and 300 mg/kg) was studied and compared with known drug compound; Furosemide. Treated animal urine was analyzed for urinary excretion and diuretic action. The anti-hypertensive effect was statistically significant when compared with the control p < 0.001. The extract at 100mg/kg demonstrated the best systolic and diastolic blood pressure lowering potential as compared to other concentrations. The diuretic action of the plant extract at the lowest dose (100 mg/kg) was high and quantitatively similar to the standard drug. The combined powdered leaf, stem and roots aqueous extract of *C. chinense* possesses anti-hypertensive and diuretic potential in salt loaded hypertensive rats.

## INTRODUCTION

Hypertension, also known as high blood pressure, is a condition in which the blood vessels have persistently raised pressure or a state in which the force of the blood against the artery walls is too high (W.H.O., 2019; M.C.M. 2019). Each time the heart beats, it pumps blood into the vessels which is then carried to all parts of the body. Blood pressure is created by the force of blood pushing against the walls of the blood vessels (arteries) as it is pumped by the heart. It is written as two numbers: the first (systolic) number represents the pressure in blood vessels when the heart beats while the second (diastolic) number represents the pressure in the vessels when the heart rests between beats.The higher the pressure, the harder the heart has to pump. An elevated systolic blood pressure above 140 mmHg or a diastolic pressure above 90mmHg results into Hypertension (William, 2018).

Diuretics cause the kidneys to remove sodium and water, they relax blood vessel walls leading to the lowering of blood pressure. Hypertension can increase the risk of heart, brain and kidney diseases. It is a major cause of premature death worldwide with over a billion people having the condition (WHO, 2019). It has been reported that by the year 2025, the global prevalence of hypertension is projected to increase to 29.2% of adult population (Kearney *et al*., 2005). Its prevalence rises with advancing age with more than half of people aged 60 to 69 years been affected (Ayub *et al*., 2010). The proportion of the global burden of ailment attributable to hypertension has significantly increased from about 4.5% (nearly1 billion adults) in 2000, to 7% in 2010 (WHO, 2008; AU, 2013). Hypertension, especially salt sensitive hypertension, is more common in the black race than in Caucasians (Luft and Weinberger, 1997; Fields *et al*., 2004; O’Shaughnessy and Karet, 2004). Its development may be associated with oxidative stress (Touyz and Briones, 2011) or can be caused by a number of risk factors which includes: excessive sodium intake, obesity, alcohol intake, caffeine intake, smoking etc. (Grundy SM, 2002; Macmahon S. 1987; Fuchs, FD *et al*., 2001 and Rainne DG, 1994).

Despite the prevalence of hypertension and its associated complications, only 29% of patients with hypertension are treated and only 45% of those given anti-hypertensive medications have been controlled. The management or treatment of this ailment however centres on lowering the systolic blood pressure below 140 mm Hg and the diastolic blood pressure below 90 mm Hg. Its treatment also involves: lifestyle modifications and drug therapy. However the use of medicinal plants and consumption of plant or fruits based diets rich in potassium has proven to be very potent at effectively managing this ailment (Bruneton J, 1995; David LH, 2005; Appel *et al*., 2006; Androgue and Madias, 2007).

*Clerodendrum chinense* (Osbeck) Mabberley (Synonym: *Clerodendrum fragrans* Willd., *Clerodendrum philippinum* Schauer) belong to family Verbenaceae. A semi woody shrub distributed in Southern Asia commonly known as Chinese glory tree, Scent malli and Brajamalli in the state of Odisha, India. It is known for its wild growth, vegetative spread and as an ornamental plant (Venkatanarasimman *et al*., 2012 and Satapathy *et al*., 2010). The use of various parts of *Clerodendrum* species as folk and traditional medicines have been reported in India, China, Korea, Japan, Thailand, Africa etc. This include: remedial purpose in inflammatory disorders, diabetes, cancers, malaria, fever, etc (Shrivastava and Patel, 2007).

*Clerodendrum chinense* is known in folklore medicine for their anti-hypertensive and diuretic properties, however little studies have been conducted to validate this proof (Nguyen Van Duong, 1998). The juice from this leaves is an ingredient of herbal bath for children with furuncles. (Infection of the hair follicle most commonly caused by the bacterium *Staphylococcus aureus*). A decoction of the leaves is reputedly a remedy for difficult cases of scabies. The plant is used in a fomentation for the treatment of rheumatism and ague (malaria or some other illness involving fever and shivering) and as an ingredient in a mixture for treating skin problems. Its root is considered antibacterial, antiphlogistic (anti-inflammatory) and diuretic. It is used in the treatment of abdominal pain, intestinal diseases, kidney dysfunctions etc. (Mihir *et al*., 2015). It is said to have been used successfully in the treatment of jaundice and a wide range of women’s disorders, skin afflictions, lumbago and hypertension owing to its diuretic properties. A decoction of the leaves is applied externally as an antiseptic. It is used for treating rheumatic arthritis, beri-beri, edema, leukorrhea, bronchitis, hemorrhoids, prolapsed rectum, scrofula, chronic osteomyelitis. The class of constituents present in this plant species are terpenes which include: monoterpenes, diterpenes, triterpenes, iridoids and sesquiterpenes (Barua and Chowdhury, 1989). Other major chemical constituents reported from *Clerodendrum chinense* are phenolics, flavonoids, terpenoids, steroids, etc. (Kar *et al*., 2015).

This present investigation deals with the evaluation of anti-hypertensive and diuretic activity of the aqueous leaf, stem and root extracts of *Clerodendrum chinense*.

## MATERIALS AND METHOD

### MATERIALS

#### Drugs and chemicals

The drug and chemical reagents used in this study include; Distilled water, normal saline, Furosemide. Other chemicals and reagents used for phytochemical tests were obtained from Department of Pharmacognosy, Faculty of Pharmacy, University of Lagos.

#### Experimental animals

Adult Female wistar rats, weighing 83–95 grams, were purchased from Animal house, Department of Pharmacology, College of Medicine, University of Lagos, Nigeria. The animals were kept in plastic cage in the animal house of the Department of Pharmacology, College of Medicine, with 12 hour/light dark cycle. The animals were allowed free access to standard laboratory pellet and tap water. Prior to the start of the experiment all animals were fasted overnight with water *ad libitum*. The care and handling were in accordance with the internationally accepted guidelines for use of experimental animals (Vogel, 2007; OECD, 2008).

#### Collection of the plant

The leaf, stem and root of *Clerodendrum chinense* were collected in April 2017. Verification of the plant identity was done at the Herbarium of the department of botany, University of Lagos, Nigeria with voucher number 6974.

### METHODS

#### Extraction of the Plant

The plant parts were air dried for two weeks, coarsely powdered and extracted by weighing 548g of the powdered samples in 9.5L of boiling water until the powder was totally submerged. It was allowed to stand at room temperature for a period of 24h with constant agitation. It was then filtered using Whatman filter paper. The filtrate was oven dried at 60°C. After drying, percentage yield was calculated (Gakunga *et al*., 2013), then reconstituted in distilled water for oral administration.

#### Qualitative Phytochemical Analysis

Phytochemical studies were conducted on the combined leaf, stem and root extract to test for the presence or absence of alkaloids, flavonoids, reducing sugars, anthraquinones (free or combined), tannins, saponins, terpenoids, phlobatanins, glycosides, cardiac glycosides, phytosterols (steroidal nucleus), protein and amino acids, fixed oil and fats using the methods described by Ajayi *et al*., 2011; Harborne and Baxter, 1993; Sofowora, 1993; Trease and Evans, 2002.

#### Grouping and dosing of animals and Anti-hypertensive evaluation

6 female wistar rats were distributed into 6 groups. The experiments were done in line with guidelines on use of animals for experiments as issued by the Physiological Society of Nigeria and Physiological Society, London. Hypertension was induced by salt-loading rats with 8% sodium chloride diet (Sofowora *et al*., 2002; Mojiminiyi *et al*., 2007). *C. chinense* extract was also administered while furosemide served as standard drug. They were treated for 8 weeks in the following groups:

- Group 1: Control (Normal diet + water)
- Group 2: Salt-loaded (8% salt diet + water)
- Group 3: Treatment with 100mg/kg extract concentration
- Group 4: Treatment with 200mg/kg extract concentration
- Group 5: Treatment with 300mg/kg extract concentration
- Group 6: Treatment with standard drug (Furosemide 10mg/kg) Note: Normal diet= 0.3% Salt

#### Measurement of Blood pressure and Heart rate

The blood pressure was determined using the model 7D Grass polygraph which is an invasive method of blood pressure determination thus two rats each were randomly picked from the control group and the salt loaded group. Blood pressure reading was taken from the carotid artery to determine if the rats were hypertensive before inducing treatment. The trachea was cannulated to improve ventilation and the carotid artery was cannulated for the measurement of blood pressure. Immediately after the cannulation of the carotid artery, 0.2 ml of heparinized saline was injected into it to prevent intravascular coagulation. The arterial cannula was then coupled to a pressure transducer (Statham P23D) which had been previously calibrated. This was in turn connected to a model 7D Grass polygraph (Grass Instruments, Quincy, MA, USA) for the recording of blood pressure and heart rate. The systolic, diastolic and pulse pressures were measured from the recordings and the mean arterial pressure (MAP) was calculated as sum of diastolic blood pressure and 1/3 pulse pressure (Adeneye *et al*., 2006; Mojiminiyi, *et al*., 2007). Heart rates were obtained by counting the arterial pulses for 15s and then multiplied by four for conversion to beats/min (Adeneye *et al*., 2006; Mojiminiyi, *et al*., 2007). The rats were anesthetized after 10weeks with a mixture of 25% urethane and 1% chloralose given intraperitoneally at a dose of 5ml/kg body weight.

#### Diuretic Activity

The same group used for antihypertensive activity was used for diuretic test. In this experiment, the untreated (8% salt diet) served as the control as the diuretic activity of the extract was determined in hypertensive rats. The normal diet group is normotensive rats.

- Group 1 consisted of animals fed with normal diet.
- Group 2: Control group were fed with 8% Salt Diet.
- Groups 3, 4 & 5 were the three (3) treatment groups and were treated with 100mg/kg, 200mg/kg and 300mg/kg doses of the extract respectively.
- Group 6 were treated with standard drug (Furosemide 10mg/kg).

Route of administration for all groups was orally using oral canula. Diuretic activity was determined following the methods used by Lahlou *et al* (2007). Each female mouse was placed in an individual metabolic cage 24 h prior to commencement of the experiment for adaptation and then fasted overnight with free access to water. Immediately after administration, the mice were individually placed in a metabolic cage provided with a wire mesh bottom and a funnel for collecting the urine. Urine was then collected and measured for total of 16h after dosing. The urine was then stored at -20°C for further analysis.

The following parameters were calculated in order to compare the effects of the doses of extract and Furosemide on urine excretion:

1. The urinary excretion independent of the animal weight was calculated as ratio of total urinary output by total liquid administered using Formula-1:

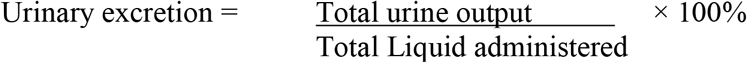
2. The diuretic action of a given dose of a drug: The ratio of urinary excretion in test group to urinary excretion in the control group using Formula 2:

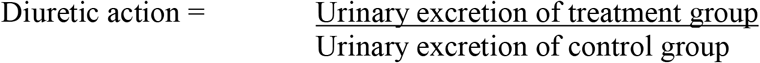
3. The diuretic activity: The diuretic action of the extracts was compared to that of the standard drug in the test group using Formula 3:

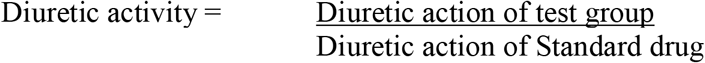

### Analytical procedures

The urine Electrolyte levels of the different animal groups were determined at FIIRO (Federal Institute of Industrial Research, Oshodi) Nigeria. Sodium & potassium level of the urine and the different extracts was determined. Na^+^ and K^+^ concentrations were measured using AAS (Atomic Absorption Spectrophotometer) AA-7000. The instrument was automatically calibrated with standard solutions containing different concentrations of sodium & potassium ions. The conductivity was measured with a Mettler Toledo conductometer on fresh urine sample.

#### Acute oral toxicity study

Acute oral toxicity test was carried out as per the Organization for Economic Co-operation and Development (OECD) guidelines for testing of chemicals number 425 (Kane *et al*., 2009 and Ancy *et al*., 2013). 6 groups (Control, Untreated, Extract at 100-300mg/kg respectively and Furosemide treated group) of 5 female wistar rats each weighing 100g-160g were fasted for 12 hour prior to the experiment and were administered with doses of 100, 200 & 300 mg/kg/day of the aqueous extract orally using oral canula. Immediately after dosing, the animals are observed continuously for the first 4 hours for physical and behavioral changes. Treatment was continued while lethality or death were also observed and recorded on Day 1, 7 and 14 respectively. The histological analysis of kidney, liver and brain was carried out at the Department of Anatomic and Molecular Pathology College of Medicine, Lagos.

### Statistical Analysis

Statistical analysis was performed. All results was expressed as means ± standard error of mean and statistical analysis was carried out by ANOVA and Bonferroni’s multiple comparison Test using Graph Pad Prism 5.0. Differences were considered significant at p <0.05.

## RESULTS

The powdered plant sample (leaves, stem and roots) at weight 548g produced an extract at weight of 26.8g. A percentage yield of 4.9% was gotten.

### Qualitative Phytochemical Analysis

Table 1 represents the preliminary phytochemical screening reports of the combined leaf, stem and root aqueous extract of *Clerodendrum chinensis*. Extracts was found to contain flavonoids, carbohydrates, tannins, saponins, terpenoids, phlobatanins, cardiac glycosides, phytosterols and proteins.

**Table 1:** Preliminary phytochemical screening of *Clerodendrum chinensis* (Osbeck) Mabberley leaf, stem and root aqueous extract.

| PHYTOCHEMICALS | Result |
| --- | --- |
| Alkaloids | - |
| Reducing Sugars (Carbohydrates) | + |
| Flavonoids | + |
| Anthraquinones | - |
| Glycosides | - |
| Tannins | + |
| Saponins | + |
| Terpenoids | + |
| Phlobatanins | + |
| Cardiac Glycosides | + |
| Phytosterols | + |
| Proteins and Amino acids | + |
| Fixed Oils and Fats | - |

‘+’ indicates present, ‘-’ indicates absent

### Effect of 8% NaCl diet in normal rats

The effect of 8% NaCl diet depicted in table 2 on randomly selected female wistar rats from the control group were hypertensive after 8 weeks. The salt loaded group indicated a high systolic and diastolic blood pressure of 130.8 ± 1.2 and 121.7 ± 1.2 respectively as compared with their blood pressure before the salt diet was introduced (Control group: 98 ± 2.6 and 75 ± 5.2 respectively). The mean arterial blood pressure, pulse pressure as well as the heart rate also increased on introducing the salt diet.

**TABLE 2:**
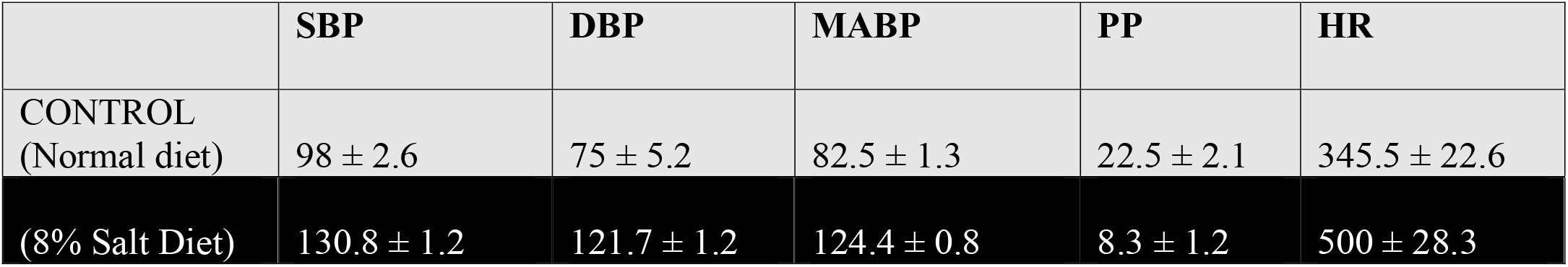
Effect of 8% NaCl diet in normal rats. SBP= Systolic Blood Pressure, DBP= Diastolic Blood Pressure, MABP= Mean Arterial Blood Pressure PP= Pulse Pressure, HR= Heart Rate.

### Effect of extracts on 8% NaCl diet loaded hypertensive rats

After 2 weeks oral administration of *C. chinense* aqueous extract, the lowest dose (100 mg/kg) of the aqueous extract showed significant decrease in mean arterial blood pressure from 143.8 ± 3.2 to 67.6 ± 5.7mmHg. In animals receiving the doses of 200 and 300 mg/kg, mean arterial blood pressure were 68.8 ± 2.4mmHg and 62.9 ± 4.7 mmHg respectively. The effect was statistically significant when compared with the control (p < 0.001) while the heart rate (HR) were statistically non-significant at 100, and 200mg/kg respectively. *C. chinense* at100 mg/kg decreased the systolic pressure to 81.3 ± 6.1mmHg and the diastolic pressure to 60.8 ± 6.9 mmHg, 200 mg/kg decreased the systolic pressure to 79.7 ± 2.3 and the diastolic pressure to 63.4 ± 2.5 mmHg while 300 mg/kg decreased the systolic pressure to 75.8 ± 5.7 and the diastolic pressure to 62.9 ± 8.4 mmHg respectively (Table 3).

**TABLE 3:** Effect of extracts on 8% NaCl diet hypertensive rats.

| TREATMENTS | SBP | DBP | MABP | PP | HR |
| --- | --- | --- | --- | --- | --- |
| Control-Normal diet | 109.4 ± 3.9 | 85 ± 9.8 | 93.4 ± 6.9 | 25.5 ± 9.3 | 340 ± 48.9 |
| 8% NaCl diet | 154.6 ± 2.4<sup>***</sup> | 133.8 ± 3.2<sup>***</sup> | 143.8 ± 3.2<sup>***</sup> | 22.7 ± 1.7 | 540 ± 45.0<sup>***</sup> |
| Extract (100mg/Kg) | 81.3 ± 6.1<sup>***</sup> | 60.8 ± 6.9<sup>***</sup> | 67.6 ± 5.7<sup>***</sup> | 20.4 ± 7.0 | 380 ± 26.2<sup>ns</sup> |
| Extract (200mg/Kg) | 79.7 ± 2.3<sup>***</sup> | 63.4 ± 2.5<sup>***</sup> | 68.8 ± 2.4<sup>***</sup> | 16.25 ± 1.3 | 307.5 ± 19.8<sup>ns</sup> |
| Extract (300mg/Kg) | 75.8 ± 5.7<sup>***</sup> | 56.3 ± 4.0<sup>***</sup> | 62.9 ± 4.7<sup>***</sup> | 20.2 ± 3.0 | 277.5 ± 29.0<sup>*</sup> |
| Furosemide (10mg/Kg) | 65.9 ± 8.4<sup>***</sup> | 46.9 ± 7.4<sup>***</sup> | 52.7 ± 7.7<sup>***</sup> | 19.1 ± 2.1 | 315 ± 26.0<sup>ns</sup> |

Values are in mmHg and expressed as MEAN ± SEM. P < 0.001<sup>\*\*\*</sup> for all doses of extract and standard (Furosemide) as compared with the Control (one-way ANOVA followed by Bonferroni's Multiple Comparison Test). SBP= Systolic Blood Pressure, DBP= Diastolic Blood Pressure, MABP= Mean Arterial Blood Pressure PP= Pulse Pressure, HR= Heart Rate. Ns= Non significant.

### Diuretic activity in hypertensive rats

In table 4, Introduction of the standard (furosemide at 10mg/kg) on the hypertensive rats showed the highest diuretic action of 11.45 while on addition of the extracts at 100 mg/kg, 200 mg/kg and 300 mg/kg concentration dose, the diuretic action exhibited 8.93%, 3.96% and 4.58% respectively. The diuretic activity of the plant extract at the lowest dose was high and quantitatively similar to that of the standard-furosemide. The diuretic activity of *C. chinense* extract was 0.78 at 100mg/kg, 0.34 at 200mg/kg and 0.40 at 300mg/kg concentration dose respectively (Fig. 1).

**TABLE 4:** Effects of extract and furosemide on urinary excretion and diuretic action on hypertensive rats.

| TREATMENT | Urinary Excretion (%) | Diuretic Action (%) |
| --- | --- | --- |
| Normal Diet (10ml/Kg Distilled water) | 92.3 | 0.24 |
| CONTROL | 380.7 | 1 |
| 100Mg/Kg extract | 3400 | 8.93 |
| 200Mg/Kg extract | 1508.8 | 3.96 |
| 300Mg/Kg extract | 1742.9 | 4.58 |
| 10Mg/Kg furosemide (standard) | 4361.1 | 11.45 |

**FIGURE 1:**
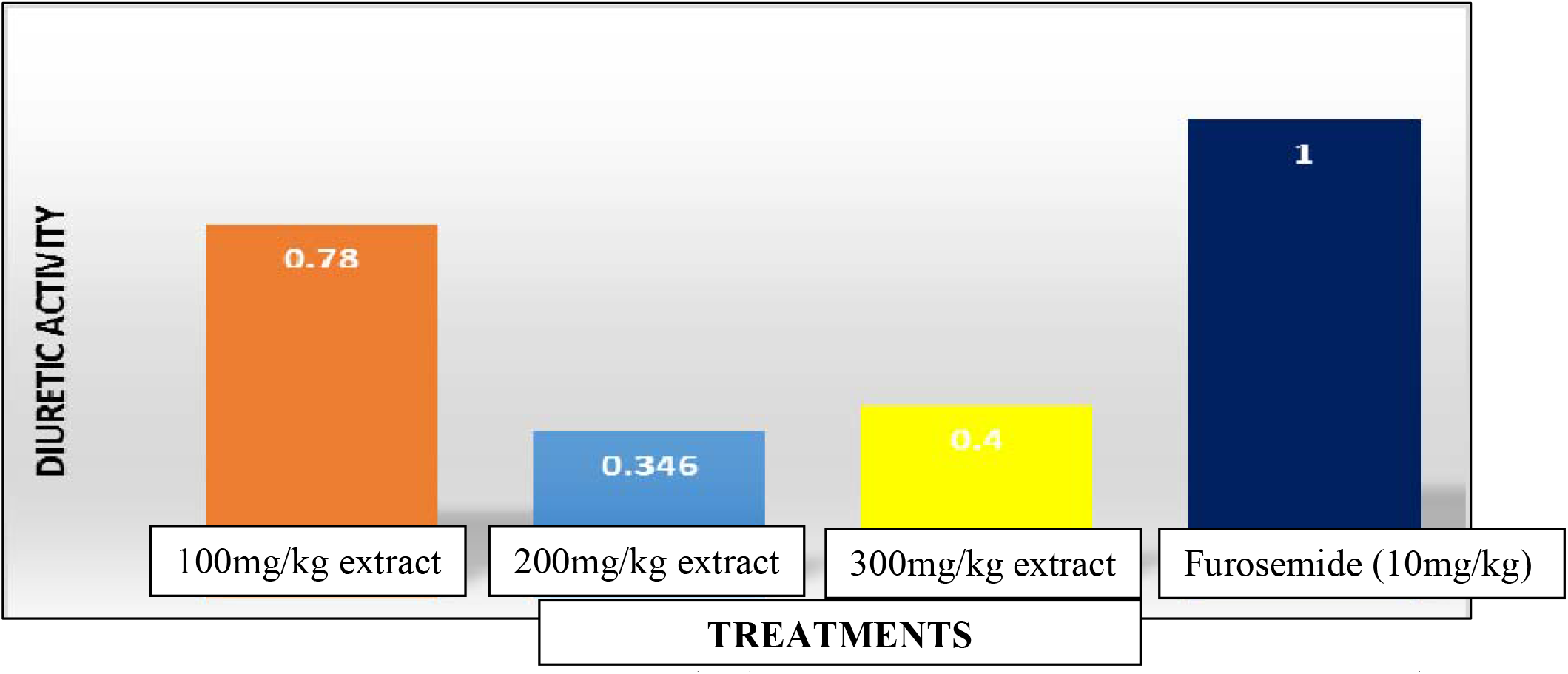
Diuretic activity of *C. chinense* extracts and Furosemide (standard) on hypertensive rats

### Effect of extract and furosemide on electrolyte concentration (Na+ and K+) in urine of hypertensive rats

Based on Na^+^ ion concentration in the urine, furosemide had the highest Na^+^ excretion (631.32 ± 471.48) followed by 100mg/kg extract (550.69 ± 197.66) while 200 & 300 mg/kg at 251.32 ± 9.39 and 374.80 ± 26.78 respectively which is lower than the control (8% Salt diet) 441.17 ± 80.96. Regarding K+ ion concentration in the urine, the control (8% salt diet) group showed more K^+^ ion excretion than the different doses of extract and furosemide (10mg/kg). The specific conductivity, which is an indirect measure of the ionic content of the urine, was increased in a dose-dependent manner in all the extract-treated groups (Table 5).

**TABLE 5:**
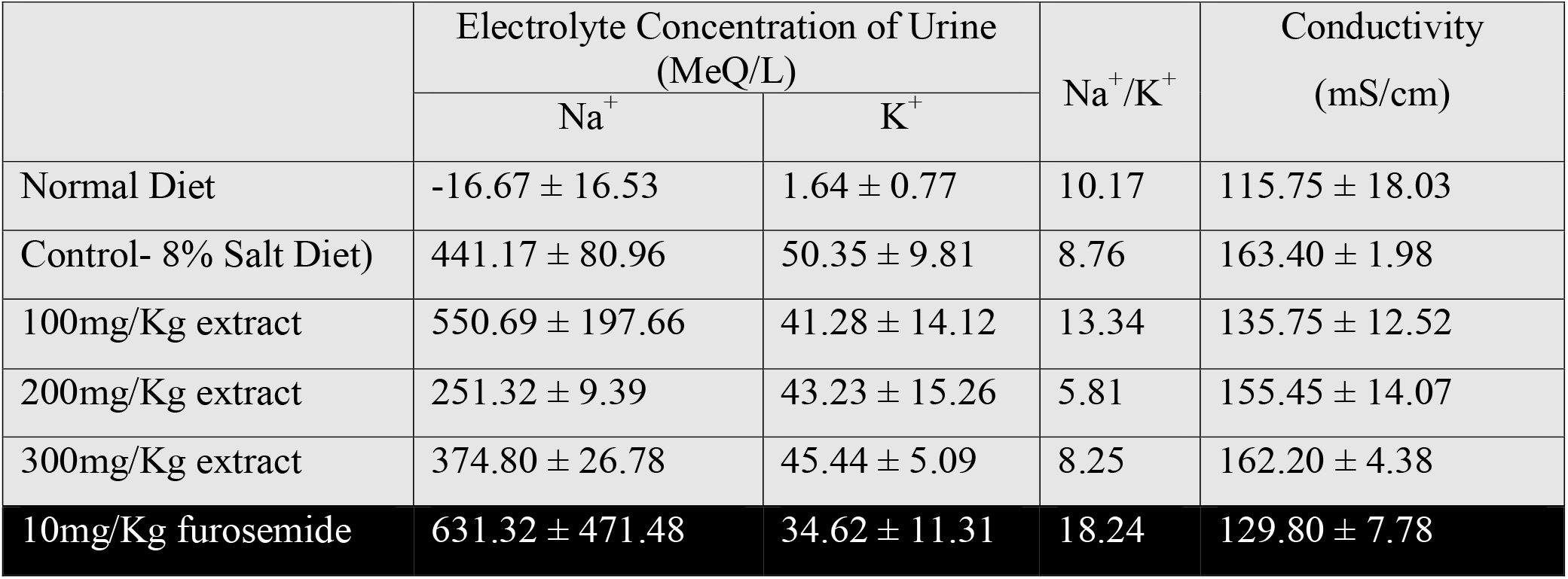
Effect of *C. chinense* extract and Furosemide on electrolyte concentration (Na+ and K+) in urine on hypertensive rats.

|  | Electrolyte Concentration of Urine (MeQ/L) |  | Na<sup>+</sup>/K<sup>+</sup> | Conductivity (mS/cm) |
| --- | --- | --- | --- | --- |
|  | Na<sup>+</sup> | K<sup>+</sup> |  |  |
| Normal Diet | -16.67 ± 16.53 | 1.64 ± 0.77 | 10.17 | 115.75 ± 18.03 |
| Control- 8% Salt Diet) | 441.17 ± 80.96 | 50.35 ± 9.81 | 8.76 | 163.40 ± 1.98 |
| 100mg/Kg extract | 550.69 ± 197.66 | 41.28 ± 14.12 | 13.34 | 135.75 ± 12.52 |
| 200mg/Kg extract | 251.32 ± 9.39 | 43.23 ± 15.26 | 5.81 | 155.45 ± 14.07 |
| 300mg/Kg extract | 374.80 ± 26.78 | 45.44 ± 5.09 | 8.25 | 162.20 ± 4.38 |
| 10mg/Kg furosemide | 631.32 ± 471.48 | 34.62 ± 11.31 | 18.24 | 129.80 ± 7.78 |

### Acute Oral Toxicity Study

It was observed that the extract of *C. chinense* did show any sign of toxicity rather there was a steady increase in the weight of the rat (Fig. 2) during the period of treatment without any observable change in skin & eyes etc. up to day 14 of the study. Unlike the rats in the high salt group, rats in the extract treated group were active and healthy in appearance reversing the weakness experienced by the rats before treatment thus showing a likely preventive effect of *C. chinense*.

**FIGURE 2:**
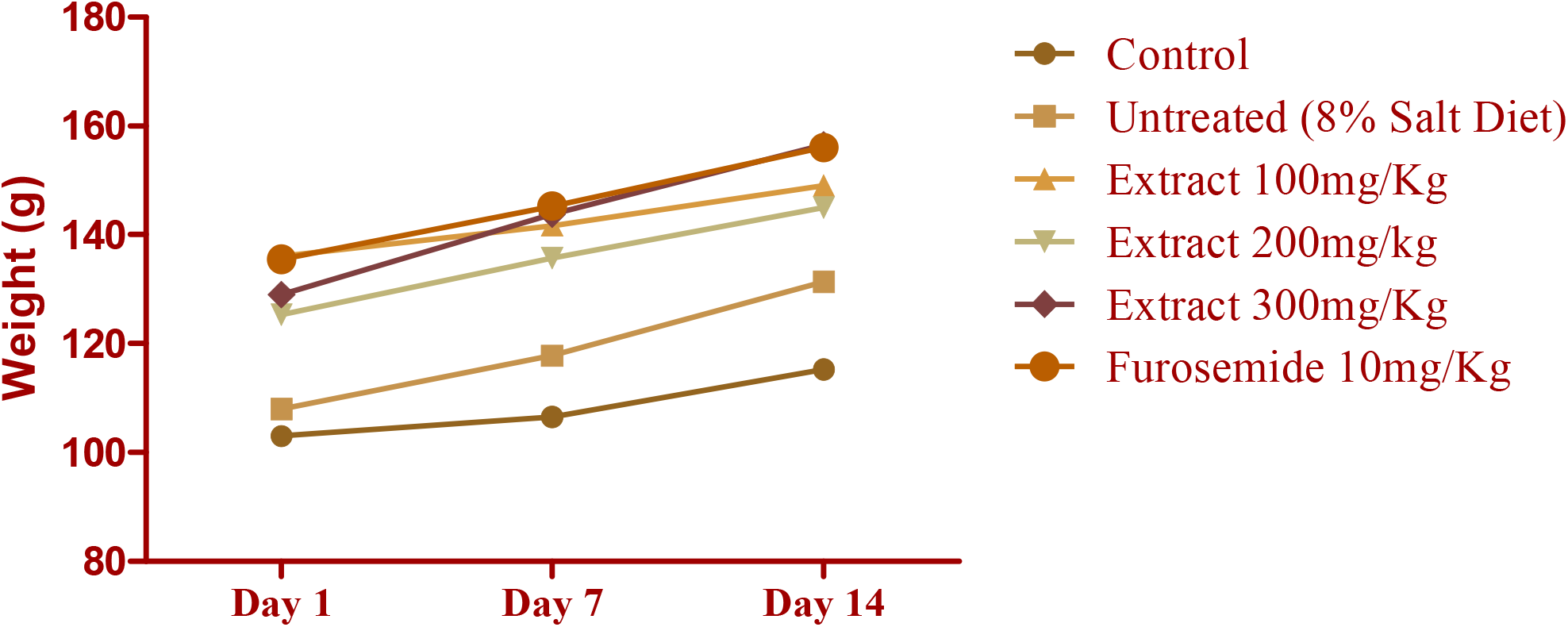
Weight change experienced by various rat groups during treatment. p < 0.0001 as compared using two-way ANOVA.

## DISCUSSION

This study investigated the antihypertensive and diuretic effects of the mixed leaves, stem and root aqueous extract of *Clerodendrum chinense* in Sodium chloride induced hypertensive rats. The preliminary phytochemical investigations in the present study revealed the presence of flavonoids, saponins and steroids (Table 1). Acute oral administration of *Clerodendrum chinense* aqueous to Sodium chloride induced hypertensive rats caused a significant decrease in systolic blood pressure, diastolic blood pressure and mean arterial blood pressure and also showed a dose dependent increase in cumulative urine volume.

Mean arterial blood pressure is the average pressure in the arteries during one cardiac cycle. The Mean Arterial Pressure is derived from Systolic Blood Pressure (SBP) and Diastolic Blood Pressure (DBP). MAP is often used as a surrogate indicator of blood flow and believed to be a better indicator of tissue perfusion than SBP as it accounts for the fact that two thirds of the cardiac cycle is spent in diastole.

Ideal and healthy blood pressure readings must be less than 120mmHg (systolic)/ 80mmHg (diastolic) but more than 90mmHg/ 60mmHg. Blood pressure readings less than 90mmHg/ 60mmHg indicated low blood pressure (Hypotension). [Blood pressure UK, 2014].

In the experiments, the female wistar rats given 8% NaCl diet were hypertensive after 8 weeks. The salt loaded group showed SBP (130.8 ± 1.3) and DBP (121 ± 1.2) compared with the control group of 98 ± 2.6 and 75 ± 5.2 respectively (Table 2). It was observed that at the 8th week of salt loading diet, the rats were trembling and seemed very weak. Much more urine was discharged, compared to the normotensive animals’ i.e Control group, so much that the bedding was often wet and had to be changed more frequently. The normotensive rats in other groups were still active and didn’t show such weakness. 6 out of 30 female wistar rats receiving 8% salt diet died within 8 weeks. This could probably be due to development of Left ventricular hypertrophy, heart failure or other serious cardiovascular diseases during the 8 weeks of salt loading (Inoko M, *et al*, 1994). The effect of 8% NaCl diet on selected wistar rats from the control group were hypertensive after 8 weeks. The salt loaded group indicated a high systolic and diastolic blood pressure as compared with their blood pressure before the salt diet was introduced (Table 2).

Readings were however taken on continuous salt loading. The systolic pressure of the rats reached 154.6 ± 2.4 mmHg and diastolic pressure reached 133.8 ± 3.2 mmHg in the untreated group (*p*<0.001 compared to value prior to salt treatment). Concurrently after 2weeks of oral administration of *C. chinense* aqueous extract, the effect was statistically significant when compared with the control (p < 0.001). *C. chinense* extract produced a significant hypotensive effect compared to the normotensive pressure after treatment of the control group: 109.4 ± 3.9 systolic and 85 ± 9.8 mmHg diastolic. The plant extract produced a dose-dependent reduction in heart rate of the rats indicating a possibility of negative chronotropic effect by the extract (Table 3). Unlike the rats in the untreated group, rats in the extract treated group were active and healthy in appearance reversing the weakness experienced by the rats before treatment thus showing preventative effect of *C. chinense*. This experiment suggests that aqueous extracts of *C. chinense* may hence exert its antihypertensive effects via the induction of diuresis or stimulation of negative chronotropic effect.

Furosemide showed the highest Diuretic action (11.45%) followed by 100 mg/kg extract (8.93%), 300 mg/kg (4.58%), 200 mg/kg (3.96%) indicating that the extract did not show a dose-dependent diuresis. The diuretic action of the plant extract at the lowest dose was high and quantitatively similar to that of Furosemide (Table 4). According to Gujral *et al*., 1955, diuretic activity is good if it is greater than 1.5, moderate if it is between 1.0 and 1.5, little if between 0.72 and 1.0 and nil if it is less than 0. Thus, 100 mg/kg aqueous extract showed little diuresis while 200 & 300mg/kg extract showed very slight diuresis compared to Furosemide (Standard) (Fig.1).

The effect of *C. chinense* extract and Furosemide on electrolyte concentration (Na+ and K+) in urine of the hypertensive rats indicated that diuretic effect is not of the natriuretic type. The control group showed more K^+^ ion excretion than the different doses of extract and Furosemide. Loop diuretics like furosemide increases urinary flow rate and urinary excretion of sodium, potassium and chloride, by inhibiting Na^+^-K^+^-2CI^-^ symporter in the Thick Ascending Loop and inhibiting Carbonic Anhydrase enzyme (Bevevino *et al*., 1994). In this experiment, the low K^+^ ion concentration in the urine compared to the control may be as a result of unknown experimental errors or the brand of Furosemide tablets used. The specific conductivity, which is an indirect measure of the ionic content of the urine, was increased in a dose-dependent manner in all the extract-treated groups (Table 5). It is however possible that *C. chinense* aqueous extract exerted diuretic effect by inhibiting tubular reabsorption of water and sodium as direct action of potassium content of *C. chinense* extract on diuretic effect is not considered, since the K^+^ content of the extract was very low in comparison with the Na^+^ concentration obtained. Only 100mg/kg *C. chinense* extract produced little natriuresis.

One can suppose that the phytochemical substances present in the plant might be responsible at least in part, for the observed diuretic activity and that they may act individually or synergistically. Previous studies have demonstrated also that there are several compounds which could be responsible for the plants diuretic effects such as flavonoids, saponins or organic acids (Maghrani *et al*., 2005). The effect may be produced by stimulating regional blood flow or initial vasodilatation, or by producing inhibition of tubular reabsorption of water and anions (Mart’in-Herrera *et al*., 2008).

## CONCLUSION

This study hereby indicates that aqueous extract prepared from the combination leaves, stem and roots of *C. chinense* has antihypertensive potential. Daily oral administration of 100, 200, 300 mg/kg can result in significant hypotension. *C. chinense* extract at a low dosage produces significant diuretic activity.

